# Unilateral corneal insult in Zebrafish results in a bilateral cell shape and identity modification, supporting wound closure

**DOI:** 10.1101/2021.03.21.436164

**Authors:** Kaisa Ikkala, Vassilis Stratoulias, Frederic Michon

## Abstract

Most of terrestrial and aquatic vertebrates are equipped with camera-type eyes, offering a focused and clear sight. This apparatus is rendered inefficient if its most superficial and transparent element, the cornea, is opaque. This structure, prone to environmental aggressions, bears excellent wound healing capabilities to preserve vision. Up to date, most of the corneal wound healing studies are made on mammals. Here, for the first time, zebrafish is used as model to study wound closure of corneal epithelium after abrasion. Our study demonstrates a swift wound closure after corneal insult. Interestingly, a unilateral wound induces a bilateral response. While cell proliferation is increased during wound closure, this parameter is not crucial, and cell rearrangements seems to be the driving force. Furthermore, we discovered a profound change in epithelial cell transcriptomic signature after abrasion, reflecting a modulation of cell identity and increase of phenotypic plasticity. The latter seems to unlock terminally differentiated cell capacities for wound healing, which could be the key for a speed up organ regeneration. Our results prove that zebrafish cornea is a powerful model to investigate, not only corneal wound healing, but ectodermal organ pathophysiology.

## Introduction

Since the first simple photosensitive cells, that can still be found in some mollusks and worms, eyes have evolved towards a complex anatomy. A direction in eye evolution was the generation of the anterior segment structures, which are needed to focus light on the photoreceptors forming the retina. One of the most complex eye types is the camera-type eye and is found both in aquatic and in terrestrial animals. The main innovation of this eye type was the generation of lens and cornea, both structures derived from the ectoderm and transparent. The refractive lens focuses precisely the light onto the retina (Ayala, 2007). The cornea is a thin structure serving two roles. First, the highly cohesive epithelial cells protect the eye inner chamber from pathogens and water loss. Then, corneal organization and transparency form a refractive layer, key element for clear sight. In terrestrial animals, corneal microenvironment is composed of cell-cell contacts, dense innervation and tear film. The latter is source of hydration and nutrients to the epithelium (Zieske, 2004). When the tear film is defected, a progressive corneal degeneration is triggered, which can lead to corneal opacification, and ultimately corneal related blindness. Evidently, in aquatic environment, the tear film is lacking, and there is no evidence on the source of nutrients for corneal epithelium.

Being the most external tissue of the eye, cornea has to withstand regular aggressions. As the ocular surface is an exposed tissue, it is regularly challenged by physical damage. Corneal abrasions are very common, and lead to an important influx of patients in hospitals (Guier and Stokkermans, 2021). They are often caused by small particles, such as dust, sand or other blowing debris, and can induce a scratch on the cornea. The resulting corneal injury causes physical and functional discomfort. The subsequent edema provokes photophobia and decreased visual acuity. In case of a deeply embedded foreign object, corneal irregularities might form, resulting in significant visual disruption. Previously, we showed that murine corneal wound closure was based on cell rearrangements (Kalha et a.l, 2018a), and tear composition was quickly modified after corneal insult (Kuony et al., 2019). These changes in corneal microenvironment, undoubtly supporting corneal healing, were crucial for proper wound closure, by bringing a new mix of factors through the tears when needed, for instance to support corneal wound healing after insult.

Fish possess camera-type eyes. Their cornea is anatomically similar to mammalian ones. An adaptation to withstand pressure can be found in corneal stroma of diving fish, such as tuna (Parravicini et al., 2012). An elegant study demonstrated that epithelial renewal is similar in zebrafish and mammalian cornea (Pan et al., 2013). Indeed, both forms of epithelial renewal were found in mouse (Kalha et al., 2018a) and zebrafish (Pan et al., 2013). A local one, based on progenitor cells, and a global one, based on stem cells located at corneal periphery in a structure called limbus. Cells produced by stem cells differentiate on their way to corneal center (Strinkovsky et al., 2021). Because of this centrepetrical global cell movement, adult cornea can be divided into three territories: limbus, peripheral and central cornea. Each territory displays specific cell molecular identity (Di Girolamo et al., 2015), and dynamics (Kalha et al., 2018a).

The main difference between aquatic animal eyes and terrestrial ones is the lack of tear film covering corneal territory. The main protection comes from the mucus layer covering the whole animal. Therefore, corneal microenvironment is based on intrinsic factors, secreted by cells residing within the cornea, as opposed to the extrinsic factors, brought by the tear film. Hence, to study corneal wound healing mechanisms or corneal defects, in aquatic animals, these elements have to be taken into consideration. It might be why fish has not been yet used as model to study corneal pathophysiologies.

Here, we characterized the early stages of corneal epithelial wound healing in zebrafish, with emphasis on cell morphology and dynamics. Our result indicated that in homeostasis, there was a clear distinction in cell area and microridge abundancy between central and peripheral cornea, reflecting the renewal dynamics of corneal epithelium. As shown in other epithelial wound healing models, cellular rearrangements were the initial driving force for wound closure. In zebrafish cornea the response was, however, remarkably fast. A transcriptomic analysis of the repairing cornea showed a downregulation of *Pax6* expression, suggesting dedifferentiation to be a key mechanism in the healing process. Furthermore, we observed a response on the contralateral cornea, and that the wound healing on contralateral cornea is faster. These results broaden the understanding of corneal wound healing across species, and contribute to the identification of potential therapeutic targets in treating corneal vision loss.

## Results

### Epithelial wound closes rapidly after corneal abrasion

To analyze the corneal wound process in zebrafish, we adapted our mechanical abrasion methodology previously used in rodents (Kalha et al., 2018b). We used an ophthalmological burr to remove the central corneal epithelium (Fig. 1A). As we showed in mouse, the mechanical removal permitted a full removal of epithelial cells at the wound site, without an obvious impact on the underlying stroma (Fig. 1B).

**Figure 1.**
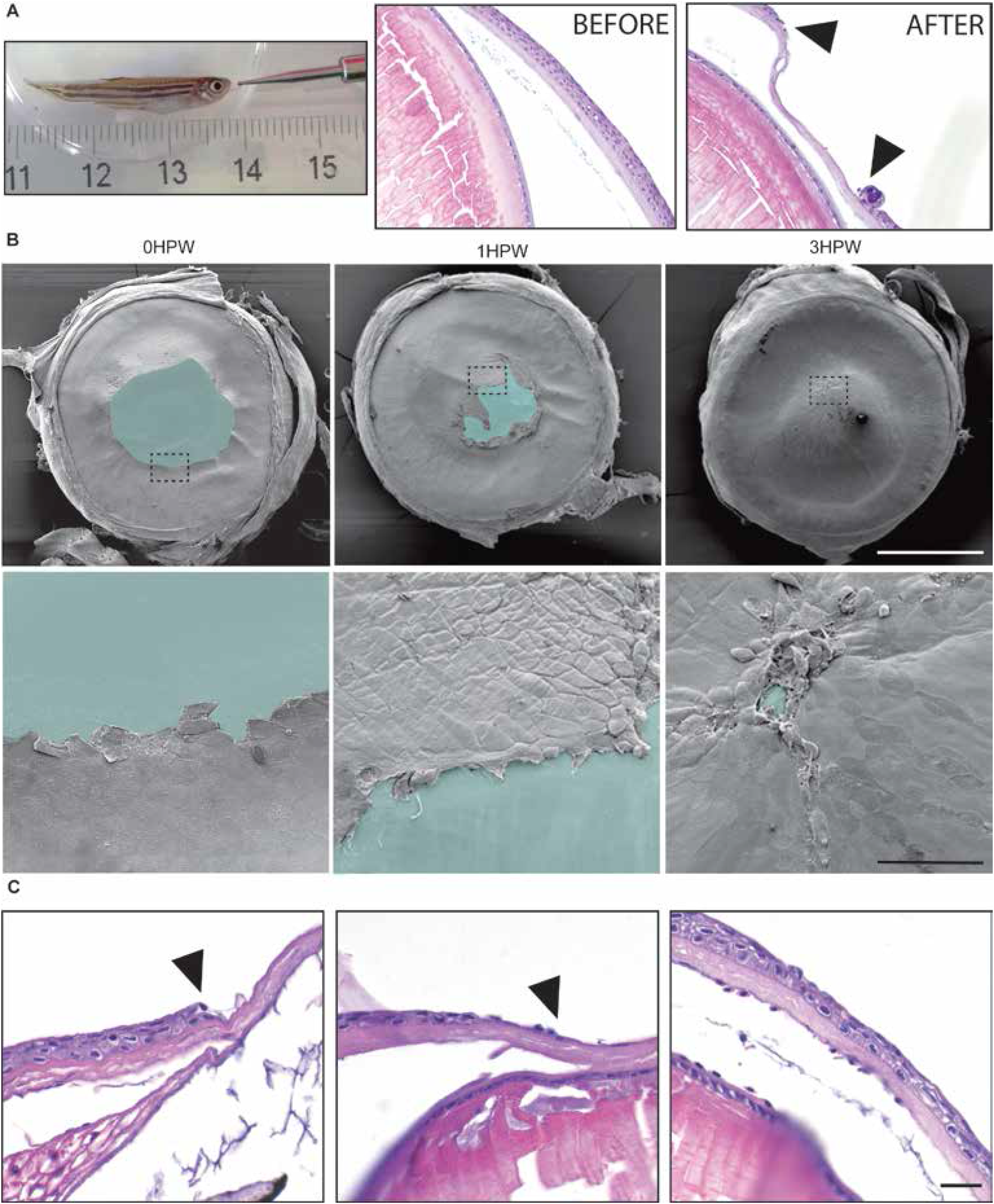
Timeframe of corneal wound closure after abrasion. **A.** Ophthalmic burr removes the epithelium from the cornea. Left: burr with 0.5 mm tip was used in this study. Right: hematoxylin-eosin staining of cornea before and right after abrasion, the arrowheads mark the limit of the wound. **B.** Scanning electron microscopy images show the wound at 0, 1, and 3 hours after abrasion. Upper panel: overview of the eyes, blue color marks the wounded area. Lower panel: close up on wound border corresponding to the dashed line boxes from upper row. **C.** Wounded area stained with Hematoxylin-Eosin shows the progressive wound closure. Scale bars: B: 500 μm upper panel, 50 μm lower panel, C: 20 μm. HPW: hours post-wound.

To gain insights about the epithelial wound healing timeframe, we used scanning electron microscopy. (Fig. 1B). Immediately after mechanical wounding, the resulting abrasion typically covered most of central cornea territory (over half of corneal radius). As little as 1 hour post-wound (1HPW), epithelial cells covered a part of the abraded area. The wound edges moved rapidly towards each other. Within 3 hours (3HPW), the wound was either completely or close to be completely sealed (Fig 1B). This timeframe was confirmed by histological sections (Fig. 1C).

To get an overview of how different regions of the cornea respond to wounding, we monitored the clonal pattern of the epithelium with the Zebrabow-M line. As reported before, this model was fitted to analyze the clonal expension during corneal epithelial renewal (Pan et al., 2013). At 6 months of age, clones extended from the periphery towards the center, reflecting limbal long-term renewal (Fig.S1A). At 3HPW, the wound was closed and all clones extended from the periphery to the wound site. By 24HPW, the clonal pattern still deviated from the pre-wound one. When comparing basal (Fig. S1B) and superficial (Fig. S1C) clonality, we saw that in physiological homeostasis the clone colors did not match, reflecting an active cell intercalation during local renewal. Interestingly, at 3HPW, the basal and superficial colors matched, hinting towards a global displacement from the periphery towards the center, and physiological homeostasis resumed within 24 hours. These observations indicated that epithelial cells from all directions participated to wound closing.

Here we convincingly showed by SEM and clonal displacement analysis that corneal closure in zebrafish is completed within 3HPW. To explain this swift reaction we investigated the mechanisms involved during corneal wound closure.

### Cell proliferation is induced after abrasion, but not required for wound closure

To understand the mechanisms in place during the corneal wound closure, we investigated the cellular events driving epithelial closure. Previously, we showed that cell proliferation is not involved in mouse for covering the wound after abrasion (Kalha et al., 2018a). To assess the requirement of epithelial cell proliferation in zebrafish during corneal wound closure, we used EdU to labelled proliferating cells (Salic and Mitchison, 2008). We added EdU to tank water for 1.5hrs (Fig. 2), whether right after corneal abrasion, or before 24HPW, by which we hypothesize that the corneal epithelium was healed, based on our previous work on murine cornea, and the time beween wound closure and healing (Kalha et al., 2018a). At 1.5HPW, the abraded cornea displayed twice as much proliferating cells as the cornea of a zebrafish that was handled the same way, without the abrasion (Fig. 2A-B). This increased proliferation was found mainly in the peripheral cornea, and more seldomly on the central cornea. Interestingly, the contralateral cornea reacted to physical abrasion. A higher number of proliferating cells was found on the contralateral side, however, this observation seemed to not be significant, due to a large variability in the response. At 24HPW, the proliferation was back to the control level, reflecting a healing phase shorter than 24 hours, and confirming our hypothesis on the wound healing period (Fig. 2C). As the proliferation seemed to be important for wound healing at the early steps of wound closure, we blocked proliferation by adding hydroxyurea in the water tank (Young and Hodas, 1964). We validated the approach by searching for EdU+ cells after abrasion (Fig. 2D-E). Hydroxyurea was a potent proliferation inhibitor. Close to no proliferating cells were found on corneas after treatment (Fig. 2E). Unexpectedly, while proliferation was induced by abrasion (Fig. 2B), inhibition of proliferation by hydoxyurea had no effect on wound closure at 3HPW (Fig. 2F).

**Figure 2.**
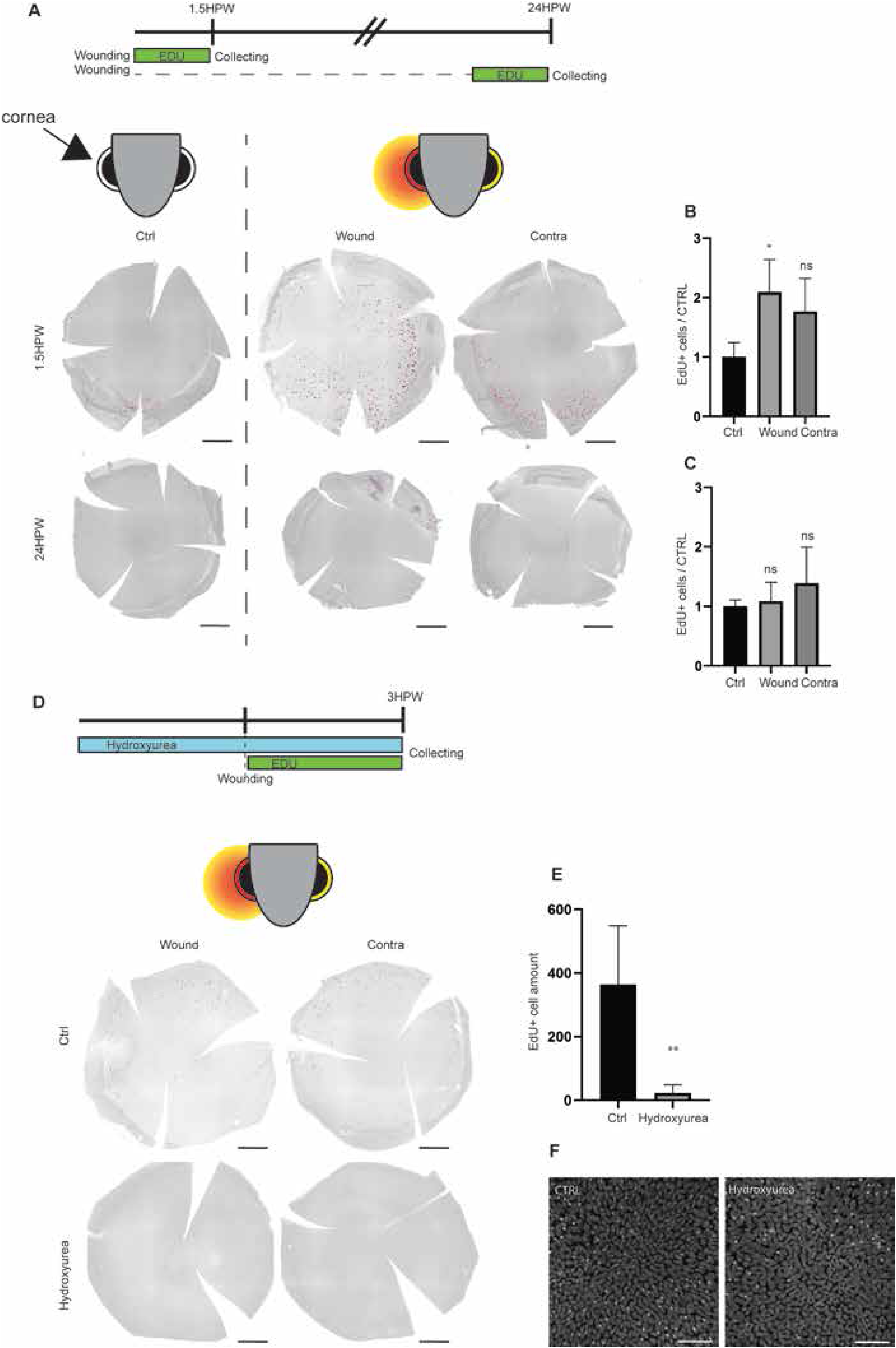
Proliferation after epithelial abrasion. **A.** EdU labeling (1.5 hours) at 1.5 or 24HPW. **B, C.** Quantification of the EdU-positive cells at 1.5HPW (**B**), or at 24HPW (C), n=3 per group. **D.** EdU labeling in animals subjected to hydroxyurea (3hrs pre- + 3 hrs after abrasion) versus controls (H_2_O). **E.** Quantification of the EdU-positive cells in **D**, n=6 per group. **F.** Hoechst staining on wound site in control versus hydroxyurea-treated animal. Quantification data represent mean + stdev, *P<0.05, **P<0.01, Kruskal-Wallis test followed by Dunn’s test for multiple comparisons, Mann-Whitney test (exact p-value, two-tailed) for comparing two groups. Scale bars: 300 μm in A and B, 50 μm in F.

### Cell morphology changes reflect rearrangements required during closure

As cell proliferation did not explain the swift wound closure, we sought for another cellular effect being responsible for covering the abraded area. In mouse, we identified cell rearrangements as the driving force for corneal wound closure after epithelial abrasion (Kalha et al., 2018). Therefore, we monitored cell shape through two parameters. Superficial cells are larger in central cornea, covering a large surface, and smaller on peripheral cornea, where most of the proliferation happens, hence, following the apical cell area should reflect changes in cellular behavior. Moreover, epithelial cell shape is well defined in cohesive epithelia, so cell roundness parameter should mirror movements. Cells were computationally analyzed for both parameters, on peripheral and central cornea, before and after abrasion (Fig. 3A). Notably, while peripheral cornea cells enlarged their apical area rapidly during wound closure (Fig. 3B), their shape did not really change (Fig. 3C). Inversely, central corneal cells, which are closer to the wound site, did not change in size (Fig. 3B), but elongated their axis toward the wound site, and therefore lost their roundness (Fig. 3C). Peripheral corneal cell shape was still affected at 24HPW, reflecting a long-term effect and longer healing than previously expected. These shape parameters reflected important modifications to close the wound.

**Figure 3.**
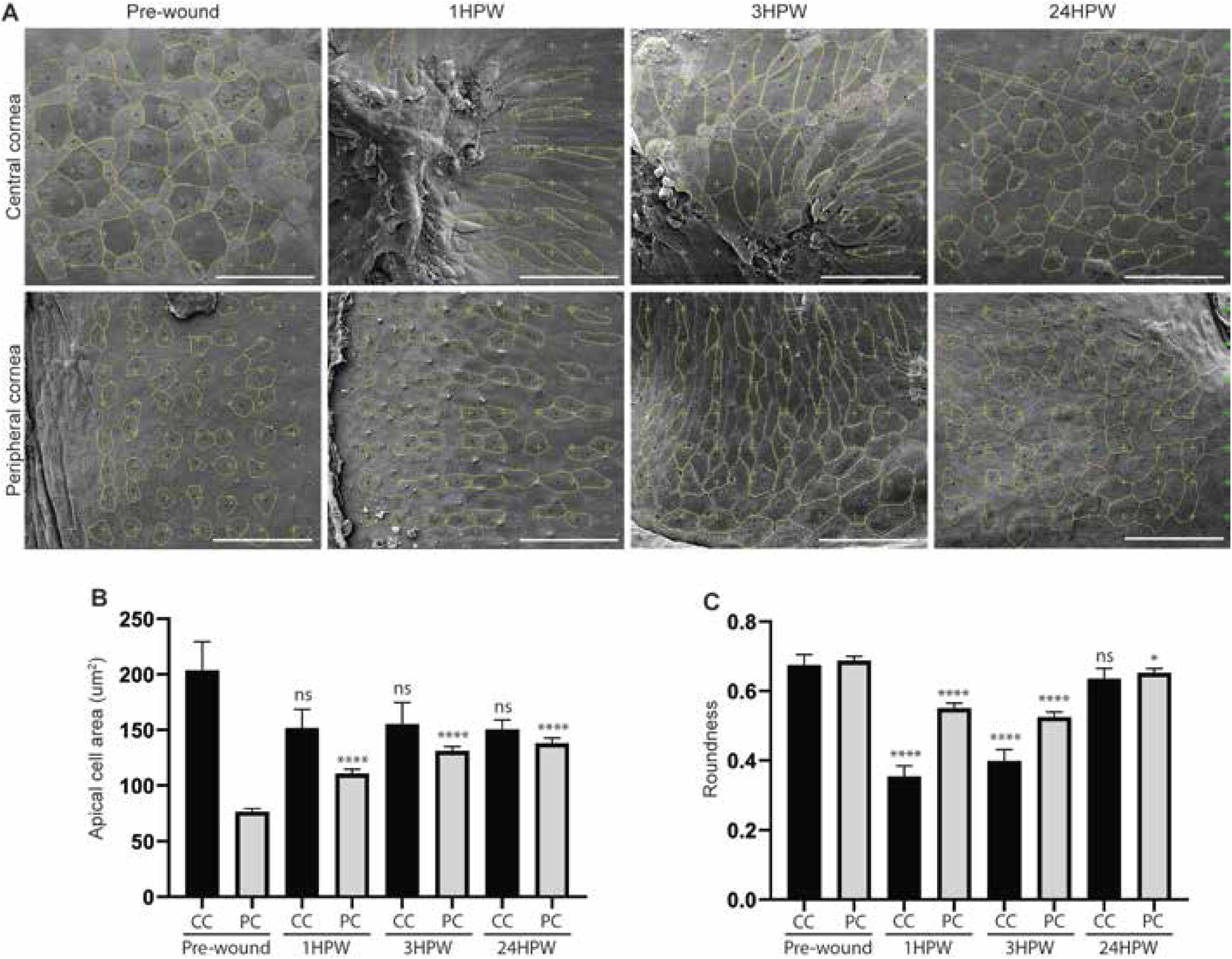
Changes in apical cell area and roundness on wounded cornea. **A.** Representative SEM images showing cell shape descriptor analysis before wounding, and 1-, 3-, and 24HPW on central and peripheral regions. **B.** Quantification of apical cell area. Cells from 3 eyes were pooled for analysis. **C.** Quantification of roundness. Cells from 3 eyes were pooled for analysis. Data represent mean+95% confidence interval, P**<0.01, P***<0.001, P***<0.0001, Kruskal-Wallis test followed by Dunn’s test for multiple comparisons. For both B and C, n=106, 85, 84, and 95 on CC, and 442, 461, 510, and 492 on PC, for pre-wound, 1HPW, 3HPW, and 24HPW, respectively. CC: central cornea, PC: peripheral cornea. Scale bars: 50 μm.

As we identified a slight effect of abrasion on contralateral corneal cell proliferation (Fig. 2B), we analyzed the same shape parameters on contralateral cornea during wound closure (Fig. 4A). Unexpectedly, despite the tight cohesion of epithelial corneal cells, peripheral corneal cells increased their apical area (Fig. 4B), and slightly elongated their axis at 3HPW (Fig. 4C). During the same period, central corneal epithelial cell did not display a significant shape modification. By 24HPW, cell shape parameters were already back to normal.

**Figure 4.**
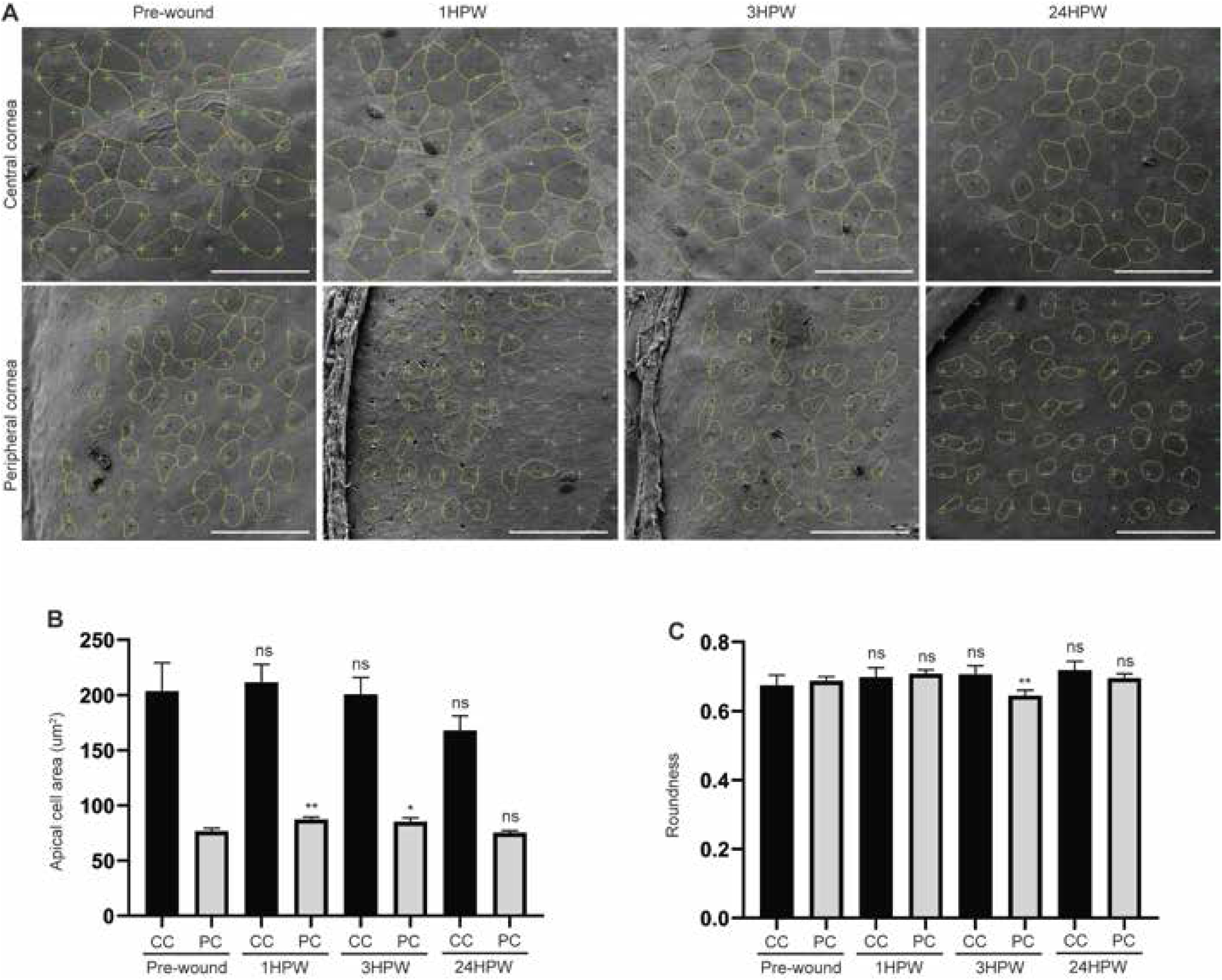
Changes in apical cell area and roundness on contralateral cornea. **A.** Representative SEM images showing cell shape descriptor analysis before wounding, and 1-, 3-, and 24HPW on central and peripheral regions. **B.** Quantification of apical cell area. Cells from 3 eyes were pooled for analysis. **C.** Quantification of roundness. Cells from 3 eyes were pooled for analysis. Data represent mean+95% confidence interval, P**<0.01, P***<0.001, P****<0.0001, Kruskal-Wallis test followed by Dunn’s test for multiple comparisons. For both B and C, n=106, 109, 129, and 72 on CC, and 442, 481, 362, and 364 on PC, for pre-wound, 1HPW, 3HPW, and 24HPW, respectively. CC: central cornea, PC: peripheral cornea, HPW: hours post-wound. Scale bars: 50 μm.

### Apical microridges are involved during wound repair

Because cell shape was affected during corneal wound closure, we investigated another element of zebrafish epithelial cell morphology, the microridges. Zebrafish, similarly to other aquatic animals, are covered with mucus, decreasing the water friction on their body. Moreover, this viscous layer offers protection from the external environment. To maintain the mucus, superficial cells possess apical protrusions, increasing their surface, and therefore their contact to the mucus layer (Pinto et al., 2019). These actin-based protrusions are subjected to a well-controlled turn-over, and an increased dynamic during wound-healing process (Lam et al., 2015). Superficial corneal epithelial cells display these protrusions (Collin and Collin, 2000). We focused on two parameters of the microridges, their total length, and their average length on each cell apical surface (Fig. 5). The first parameter describes how the cell surface is increased by the microridges, while the second parameter reflects how stable the microridges are. We discovered that peripheral corneal cells presented well defined and stabilized microridges compared to central corneal cells (Fig. 5A). To rule out the possibility that cell size could explain this phenomenon, we normalized the data to cell area. For both parameters, microridges total length (Fig. 5B) and average length (Fig. 5C), there is a significant difference between central and peripheral corneal cells. We analyzed a possible correlation between these parameters and cell area (Fig. 5D-G). Interestingly, we demonstrated that while microridge total length was mostly dependent on cell size, in central and peripheral cells (Fig. 5D-E), their average length was not (Fig. 5F-G). Meaning that microridge stability was specific to the type of cells (peripheral or central) and not on cell size.

**Figure 5.**
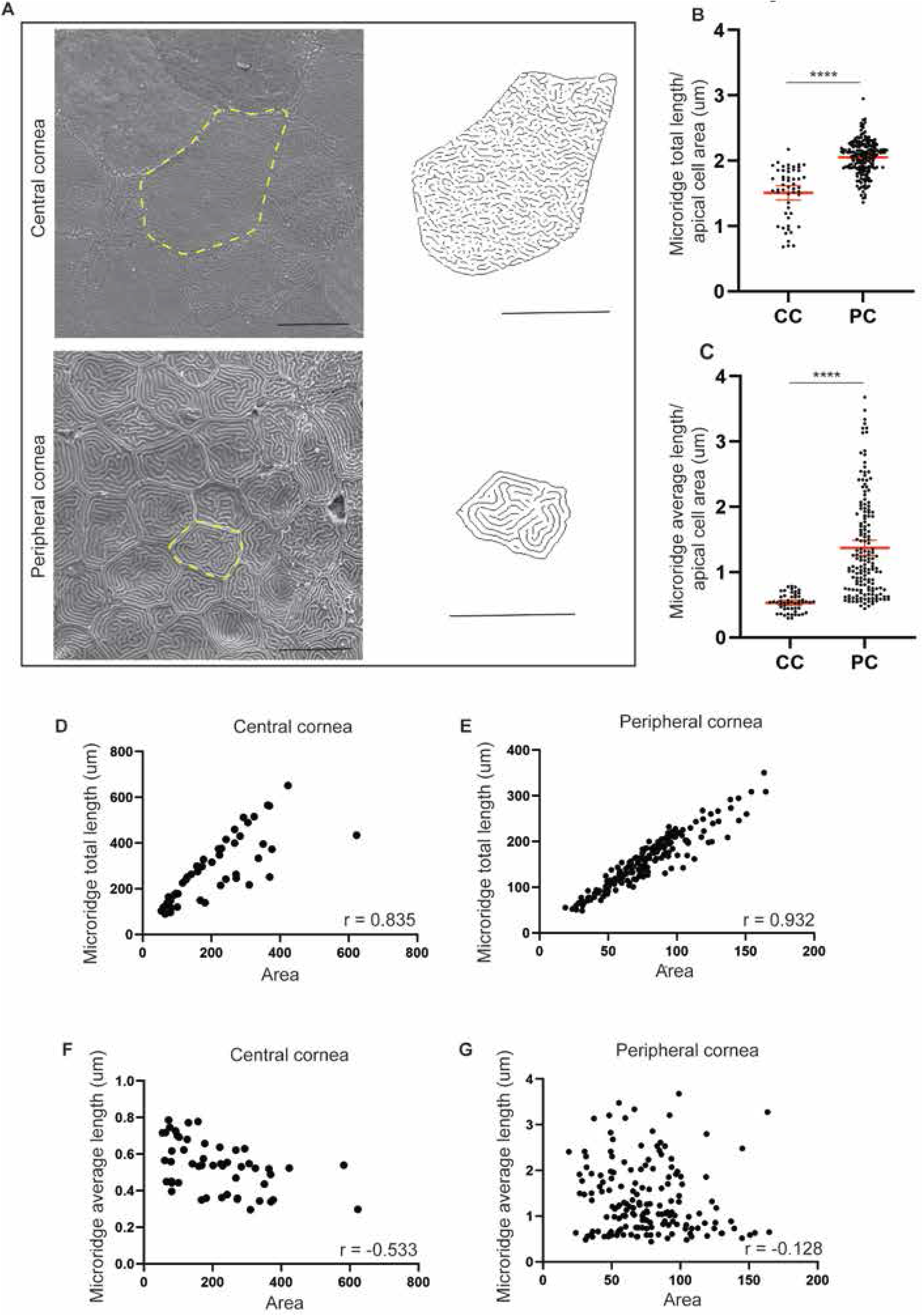
Microridges on zebrafish cornea. **A.** Representative SEM images on central and peripheral regions. **B**, **C**. Average microridge amount (**B**), and length (**C**), per apical cell area on central and peripheral cornea (n=52 for CC in **B** and **C**, n=192 for PC in **B**, and n=162 for **PC** in C after removal of outliers). Data represent mean+95% confidence interval, p****<0.0001, Man-Whitney test, two-tailed, with exact p-value. D—G. Correlation of microridge amount or average microridge length, and apical cell area, on central and peripheral cornea (n= 50 in D, 51 in E, 192 in F, and 162 in G, after removal of outliers). Spearman correlation, two-tailed, with approximate p-value, p****<0.0001. Cells from 3 eyes were pooled for every analyses. Scale bars: 10 um.

Therefore, we examined the changes in microridge dynamic during wound closure (Fig. 6A), analyzing the correlation value of microridge average length and cell size. When negative, this correlation value reflects a higher cell area and lower microridge average length, meaning high turn-over. When positive, the microridges are more stable. Our correlation analysis demonstrated that during wound closure, central corneal cells displayed a progressive increase in microridge stability, while peripheral corneal cell showed a transitory destabilization of the microridges, rapidly followed by an increased stabilization (Fig. 6B).

**Figure 6.**
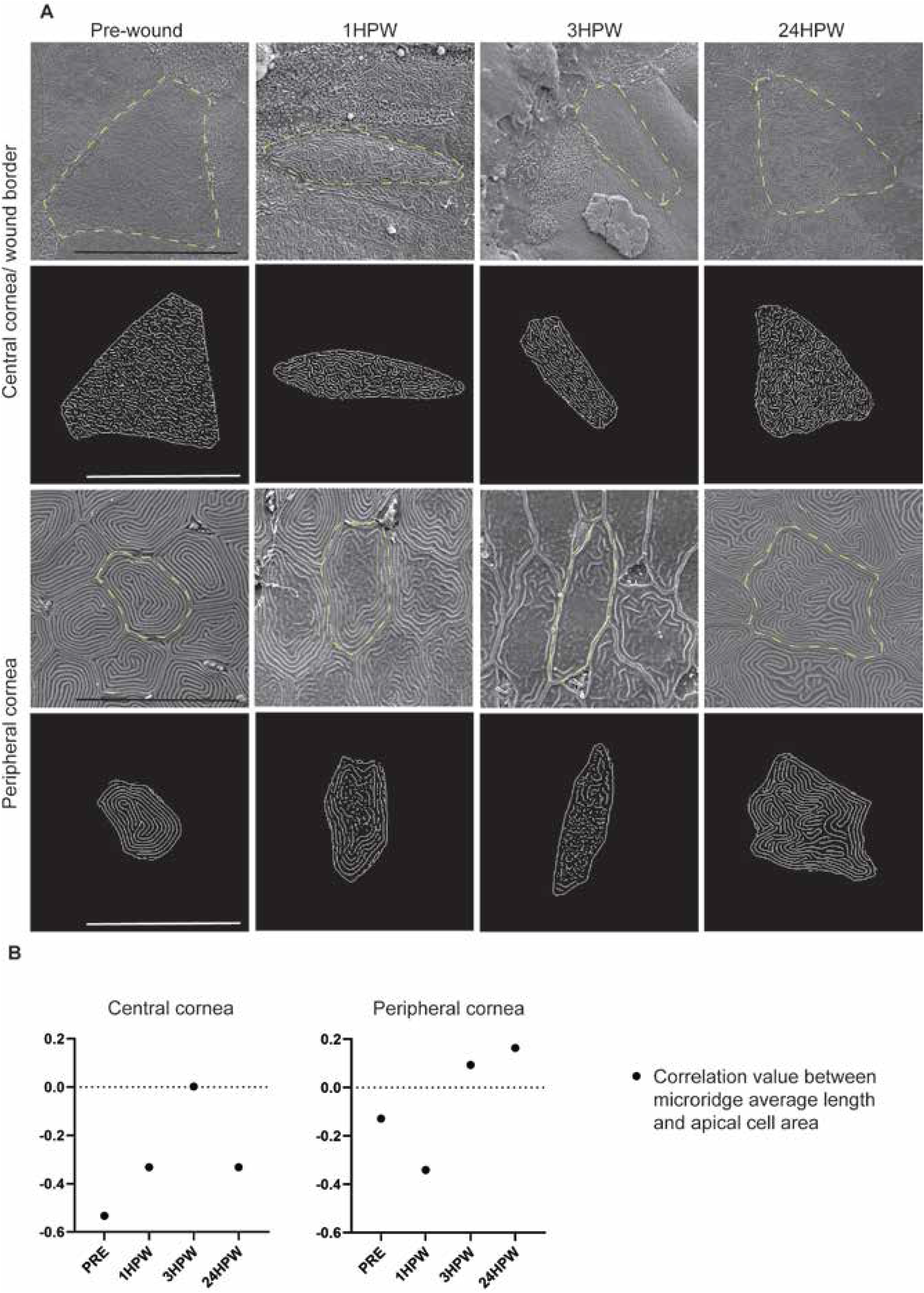
Microridges during wound closing. **A.** Representative SEM images and corresponding microridge patterns after image processing before and after wounding on central and peripheral cornea. **B.** Spearman correlation between average microridge length, and apical cell area, on central and peripheral cornea (n-numbers after removal of outliers in pre-, 1, 3, and 24HPW, respectively: CC, microridge average length: 51, 47, 30, and 43, PC, microridge average length: 162, 171, 209, and 203). Data represent mean+95% confidence interval, P**<0.01, P***<0.001, P****<0.0001. B: Kruskal-Wallis test followed by Dunn’s test for multiple comparisons. Cells from 3 eyes were pooled for all analyses. CC: central cornea, PC: peripheral cornea. Scale bars: 20 um.

### Analysis of transcriptomic changes pinpoints a bilateral epithelial cell identity modification

Because of the rapid induction of cell proliferation and shape modification, along with actin cytoskeleton turn-over alteration, we investigated modulations of transcriptomic signature during corneal wound healing.We analyzed the zebrafish corneal transcriptomic signature during wound closure, at 1.5HPW (Fig. 7A). A comparison of control *versus* wounded corneas, identified 268 genes being up- and 217 genes being down-regulated by a fold equal to or greater than 2 (Fig. 7C). We further validated 15 of these genes with qPCR (Fig. 7D). Notably, a number of genes related to eye and visual system development were significantly downregulated in wounded corneas (Fig 7B and Fig. S2). These include genes expressing the transcription factors *Pax6a*, *Hmx1*, *Six3a* and *Otx2a*, as well as *cyp1b1* a gene important for zebrafish (Williams et al., 2017) and human (Li et al., 2011) ocular physiology. Regarding *Pax6*, in Zebrafish, only *Pax6a* plays a role in ocular domain, while *Pax6b* is mainly found in the pancreas (Kleinjan et al., 2008).

**Figure 7.**
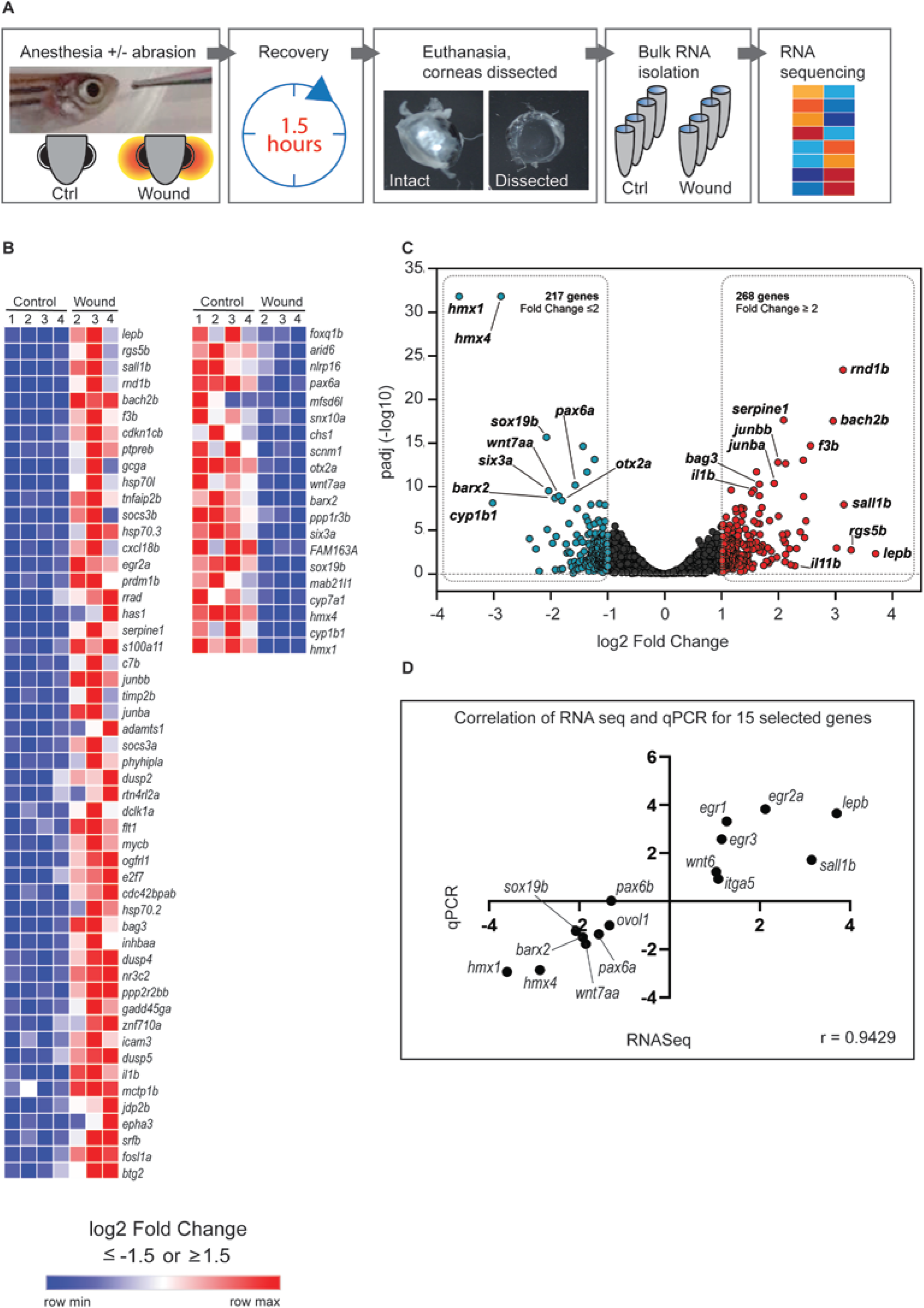
Transcriptomic changes in wound response. **A**. Schematic presentation of the experiment and sample processing for RNA sequencing **B.** Heat map of 52 most upregulated and 20 most downregulated genes (Log2 fold change ≥1.5 and ≤ −1.5, p-adjusted ≤0.05). **C.** Volcano plot of gene expression changes (wounded vs. control samples). **D.** Pearson correlation of RNA sequencing and qPCR results for selected 15 genes.

PAX6 transcription factor is known to be the master gene of eye formation (for review, Shaham et al., 2012). Previous studies reported that in the context of *Pax6* mutation, corneal cells were unable to differentiate properly, and adopt the corneal epithelial cell identity (Roux et al., 2018). After *Pax6a* expression was validated by qPCR in control and wounded corneas (Fig. 7D), we used immunohistochemistry to analyze PAX6 pattern during corneal wound healing. Before abrasion, PAX6 was found in all areas of corneal epithelium (Fig. 8A). At 1.5HPW, we detected high levels of PAX6 immunoreactivity around the wound, while the rest of corneal epithelium was void of PAX6 signal (Fig. 8B). By 3HPW, only few cells, orientated toward the wound site were PAX6 positive, while most of the cornea did not display a PAX6 signal (Fig. 8C). Notably, a strong decrease of PAX6 signal happened progressively during corneal wound healing in the contralateral eye, to the point that only few cells became PAX6 positive by 3HPW (Fig. 8F, arrows), reflecting unequivocally a bilateral response to unilateral corneal abrasion.

**Figure 8.**
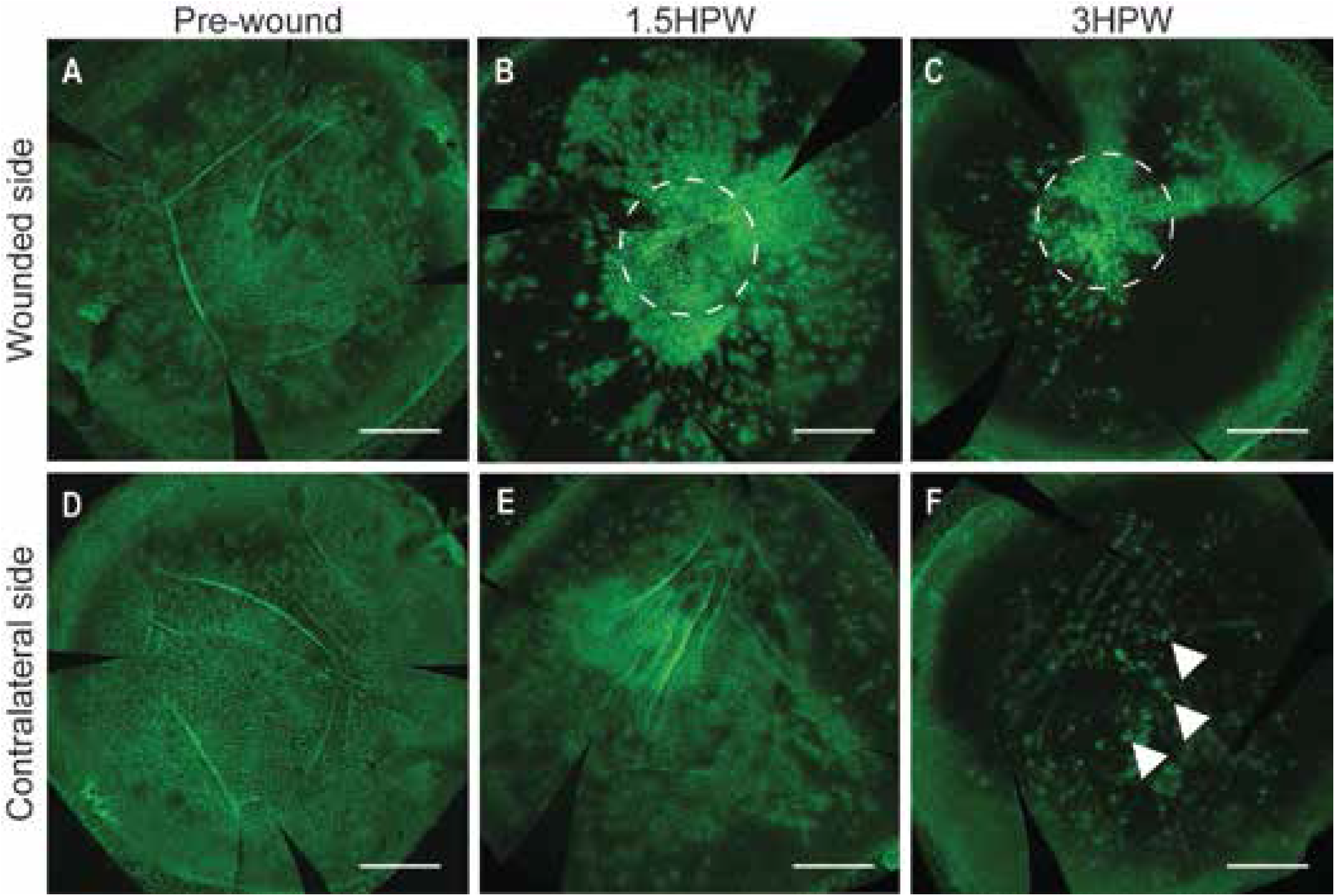
PAX6 immunological detection during wound healing. Upper panel, wounded side. Lower panel, contralateral side. Dashed lines indicate wound location. HPW: hours post-wound. Maximum intensity projection of the whole cornea, scale bars: 300 μm.

### Bilateral response prepares a swift epithelial wound healing

Finding strong cellular and molecular responses from an unharmed epithelium, which is still cohesive, was unexpected and unique. Therefore, we hypothesized that a bilateralization of corneal insult response might be beneficial in case of a secondary insult on the contralateral eye, which is still providing a proper vision. We chose to inflict an abrasion on the contralateral eye, 1 hour after abrading the ipsilateral eye, and monitor wound closure speed 1 hour after each wound (Fig. 9A). Strikingly, wound closure speed was significantly faster on the contralateral eye. While 22% of the wound was still open at 1HPW on the ipsilateral cornea, less than 5% of the wound was left to close on the contralateral eye.

**Figure 9.**
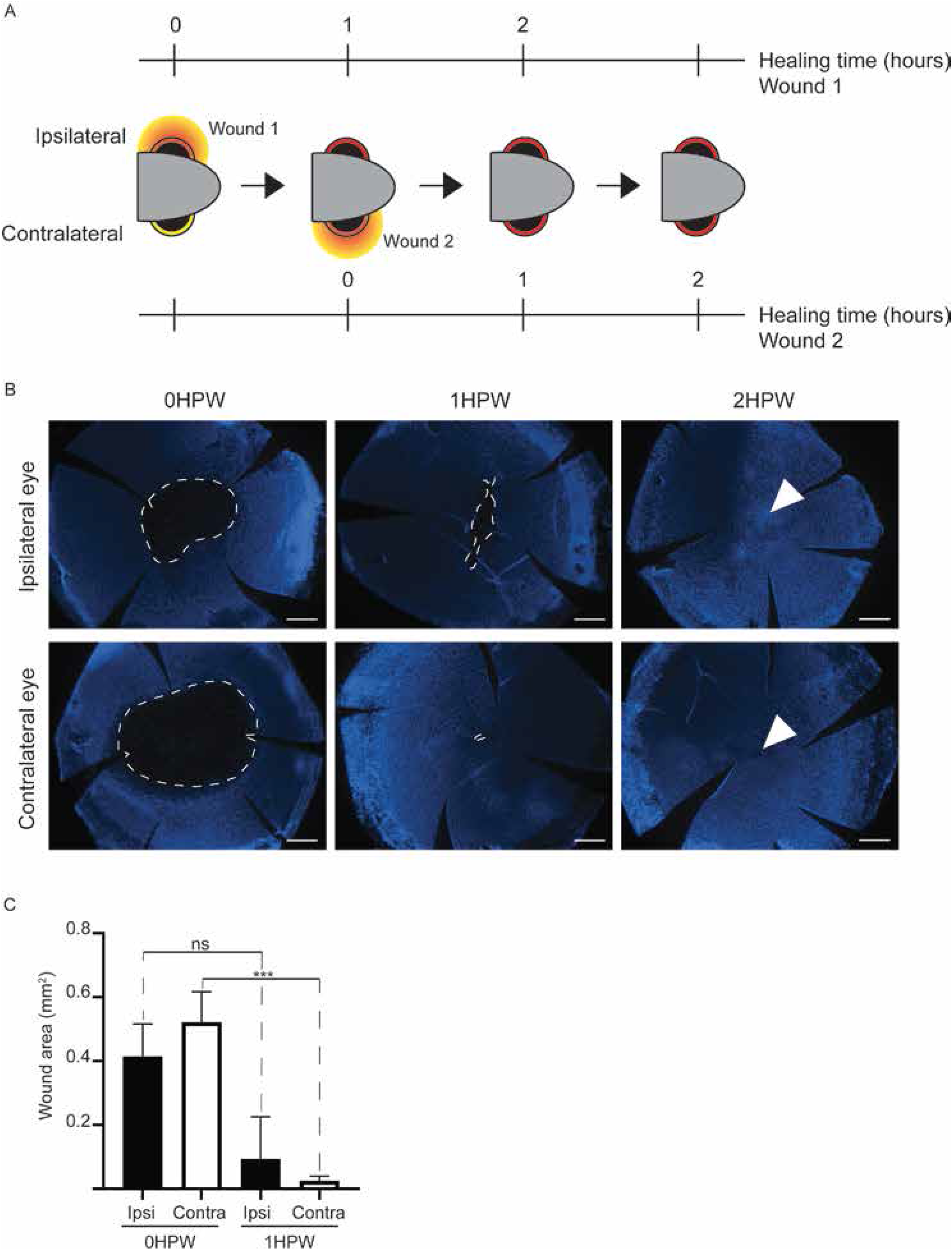
Contralateral wound healing speed. **A.** A schematic presentation of the experiment. **B.** Representative images of the cornea of either the left or the right eye at 0, 1, and 2HPW (Hoechst-staining). Dashed line indicates wound border, arrowhead points to wound site in a closed wound. **C**. Quantification of the wound area on the right and on the left side at 1 or 2HPW. Data represent mean+stdev, P***<0.001, n=5—6 per group. Kruskal-Wallis test followed by Dunn’s test for multiple comparisons. HPW: hours post-wound. Scale bars: 200 μm.

Collectively, our results confirm the bilateralization of corneal wound healing response, but more importantly point towards communication between both eyes.

## Discussion

Despite the numerous advantages of zebrafish as a model organism, especially for regenerative and wound healing processes (Sehring et al., 2016), its use in the context of corneal defects has remained unexplored. In this study, we delineated the events taking place after corneal insult in zebrafish, focusing on the cellular and molecular aspects of corneal wound closure, to compare this model to mammalian ones, and apprehend if zebrafish cornea is suitable to investigate corneal defects.

We and others have previously reported that corneal wound closure occurs in a mater of days (72 hours (Kalha et al., 2018a) and 21-55 hours (Pfister, 1975, Crosson et al., 1986) respectively). Moreover, this phenomenon did not lead to cell proliferation induction in mouse (Kalha et al., 2018a). Interestingly, our data suggest that unlike in mouse, cell proliferation was induced early on during wound closure in zebrafish. This unexpected result was confirmed by the general displacement of clones observed with the Zebrabow model. We hypothesize that this inter-species difference could be attributed to a number of factors. First, corneal wound healing in mouse is a long process. The wound closing alone takes about 3 days. The lack of reported proliferation could reflect a short proliferation window which was not picked up in previous experiments. Then, corneal wound closure might not follow the same steps in mouse and zebrafish, due to innovations brought by visual apparatus evolution. Finally, the pool of available cells for wound closure is more restricted in zebrafish compared to mouse. The lack of starting material requires then a peripheral proliferation to bring closer the wound borders. By blocking the cell proliferation induction, we ruled out the last option. Therefore, more investigation is needed to decipher thoroughly the early steps during murine corneal closure to identify the possible discrepencies between the mammalian and fish models.

Epithelial cells within a cohesive tissue are robustly stable to insure cell-cell communication, and barrier function. Our results showed an impact of the insult on cell shape (roundness) in both central and peripheral cornea, during wound healing. Cell shape modification was correlated with microridge dynamic modulation. While microridge dynamic in central corneal cells seemed to be back to normal quickly, peripheral corneal cell displayed a long term modification of microridge dynamics. These observations, together with the global epithelium displacement seen in Zebrabow model, would indicate that the driving force for wound closure is the peripheral cornea, and cells located more centrally adopt a phenotype facilitating the displacement, but are back to their original phenotype quickly after wound closure. On peripheral cornea, the cell behavior phenotype is disturbed longer. The coordination of these two territories is probably crucial for wound closure, and therefore requires specific cues to synchronize all corneal epithelial cells to achieve a swift healing.

Our transcriptomic analysis demonstrated a large modification of epithelial cell identity, especially through *Pax6* downregulation. This transcription factor is a known master gene for ocular structures, especially cornea. When *Pax6* expression is missing during eye formation in chick embryo, corneal identity fails to set in epithelial cells, and only skin identity is adopted (Collomb et al., 2013). Furthermore, a recent study demonstrated that Pax6 is an crucial player for direct cell reprogramming into corneal epithelium (Kitazawa et al., 2019). Consequently, Pax6 levels are of utmost importance when enforcing corneal epithelium identity. Our results clearly showed a specific *Pax6* downregulation. We hypothesize that loosing *Pax6* expression could lead to an increase of cell plasticity, through a less strict corneal epithelial identity. Another report showed that terminally differentiated corneal cells can adopt stem cell fate to replace lost stem cells (Nasser et al., 2018). Our original finding, together with these studies, indicate that upon injury, cell identity might be lessen to permit a swift phenotype modification, as needed during wound closure. These observations can be viewed in the larger perspective of cell plasticity in organ regeneration. By reaching a higher degree of plasticity, terminally differentiated cells seem to develop the abilities to heal a tissue or an organ, as seen in skin (Tang H et al., 2020), intestine (Kurokawa et al., 2020), and vascular system (Tombor et al., 2021), among others. Further investigations could shed new light on the molecular modulations of cell identity and phenotype required during wound healing and tissue regeneration, not only in cornea, but in most organs. It might then be possible, by selectively modify cell microenvironment, to induce enough plasticity in order to harness the regenerative capabilities.

Our previous work demonstrated the bilateralization of lacrimal gland response to unilateral corneal abrasion in mouse (Kuony et al., 2019). Unexpectedly, this response coordination on both sides seemed to play an important role as it is evolutionary conserved. The cues that are able to modify cellular and molecular behavior of a differentiated and highly cohesive epithelial tissue, without it being harmed, should be very strong. Our results pointed out not only morphological changes, and an increased proliferation, but as well a modified cell identity, through *Pax6* downregulation. Hence, our observations raise the question on the role of displaying wound healing phenotype to the contralateral cornea. At this stage, we can only speculate on the possible advantage of such mechanism. Sight is crucial for most aspects of life, such as mating, survival (as prey or predator). Therefore, loosing sight decrease chances of survival. By bilateralizing healing process, and speeding up recovering in case of bilateral injury, this mechanism increases chances of survival. While our hypothesis can only be speculative, our observation demonstrate that external cues can modify corneal epithelial identity, and these cues could help speeding up corneal would healing, which could be in the long run beneficial for patients suffering from corneal pathophysiologies.

Collectively, our results demonstrate a complex process involved in early corneal wound healing. During wound closure, not only cell rearrangements and morphology changes are involved, but as well induction of cell proliferation supports global cell displacement towards the wound site. Moreover, profound transcriptomic changes are involved rapidely after corneal insults. Together, these mechanisms are responsible for a swift wound closure, resembling to what can be observed in mammalian corneal wound healing. Therefore, we are confident that using zebrafish as corneal regeneration model would unravel new aspects of corneal physiopathologies, which could be beneficial for the 28 million people worldwide suffering from uni- or bi-lateral corneal blindness. Moreover, it would enlarge our knowledge on the connection between cell plasticity and organ regeneration, which would be beneficial not only in ophthalmology, but in all fields.

## Methods

### Fish lines and fish maintenance

Fish were maintained in standard conditions with 14h:10h light-dark cycle at the Zebrafish unit (HiLIFE, University of Helsinki). During maintenance and wound healing experiments, fish were fed one to three times a day.

For the majority of the experiments in this study we used the WT AB background (acquired from the Zebrafish facility, University of Helsinki). For analyzing the clonal pattern of the epithelial cells, we used the Zebrabow-M line ((Pan et al, 2013) a kind gift from Ivonne Sehring) crossed with a conditionally Cre-expressing line Tud104Tg ((Hans et al, 2011) Tg(hsp70l:mCherry,Cre-ERT2)tud104(AB/TL), acquired from Karlsruhe Institute of Tehnology).

Fish were collected at the age of four to seven months. The fish were randomly selected for each experiment from the stock tank.

### Corneal wounding

The fish were anesthetized with 0.02% Tricaine (A5040 Sigma) and placed into an incision on a moist sponge, head protruding from the sponge surface. Corneal epithelium wounding was performed as in (Kalha et al., 2018a/b, Kuony et al., 2019) using an ophthalmic burr (Algerbrush II) with 0.5 mm diameter tip and applying moderate pressure on the eye surface. After abrasion, the fish were transferred to tank water for recovery. Control animals were anesthetized and placed onto sponge for an equal duration as wounded animals.

### Zebrabow experimentation

The embryos from Tud104 x Zebrabow crossing were collected in E3 buffer, and subjected to Cre activation approximately 6 hours post-fertilization. 50 embryos were transferred into 1 ml of E3, and kept at +37°C for 30 minutes for inducing the expression of the recombinase protein. The tube was then kept at 28.5°C for 20—30 minutes for cooling down, and the embryos were transferred to a dish with 30 ml of 10 um 4-hydroxytamoxifen (H6278, Sigma) in E3 for nuclear translocation of the recombinase. After an overnight treatment, tamoxifen was removed by rinsing the embryos three times with fresh E3 buffer. At five days post-fertilization, the YFP-positive larvae were selected. Adult fish of four to seven months were used in wound experiments.

Wounding was done as described above. Samples were fixed in 4% PFA/PBS for 20 minutes on ice, rinsed in PBS, and stored overnight in 100% methanol at −20°C to facilitate dissecting. On the following day, the samples were permeabilized in decreasing methanol series at room temperature, permeabilized in 0.3% Triton-x-100(Q2964, MP Biomedicals)/PBS for 20 minutes, and stained with Hoechst (1:2000 in PBS, 20 minutes at room temperature). Samples were then rinsed, cornea dissected, and mounted in 80% glycerol.

### Edu-labeling, hydroxyurea treatment

For labelling proliferative cells, fish were transferred to 0.2mM Edu (900584, Sigma) for 1.5 hours before collecting samples. Three individuals were treated in the minimum of 120 ml of freshly prepared Edu solution.

In hydroxyurea treatment, fish were preincubated in 50 mM hydroxyurea (400046, Sigma) solution for 3 hours, anesthetized (0.02% MS-222), and subjected to corneal abrasion, and kept in the hydroxyurea solution for 3 hours before collecting samples.

### Whole mount staining

Fish were euthanized at selected time points by anesthesia in 0.02% Tricaine solution followed by decapitation. For whole mount staining, whole heads or enucleated eyes were fixed for 20 minutes on ice in 4% PFA prepared in PBS from 20% stock solution (15713, Electron Microscopy Sciences). The samples were rinsed with PBS and stored in 100% methanol at −20°C. The samples were rehydrated by incubation at RT in 75% methanol/25% PBS, 50% methanol/50% PBS, and 25% methanol/75% PBS, 5—10 minutes each. For samples collected as whole heads, eyes were enucleated after this step in PBS. Tissue was permeabilized with 0.3% Triton-x-100 / PBS, with rocking agitation 4 times 5 minutes RT, and blocked RT several hours (10% normal goat serum (16210064, Life Technologies), 0.5% BSA (A2153, Sigma), in 0.1% Triton-x-100/PBS) with rocking agitation. Samples were incubated with primary antibody (rabbit polyclonal antibody to Pax6, ab5790, Abcam) 1:200 in blocking solution over night at 4°C with rocking agitation. Next, samples were washed in 0.1% Triton-x-100/PBS (fast rinse, 5 times 5 minutes, 3 times 20 minutes), and blocked in blocking solution 1—2 hours at RT with rocking agitation. Samples were then incubated in secondary antibody (goat polyclonal antibody to rabbit IgG, A11008, Life Technologies) 1:200 in blocking solution, containing Hoechst 1:2000 (H3570, Invitrogen) 2—3 hours at RT with rocking agitation. Finally, samples were washed as above and stored in PBS at 4°C until dissecting. Cornea was dissected from the eye in PBS using fine scissors and tweezers and placed onto microscopy slide in a drop of 80% glycerol, and covered with a coverslip.

EdU-labeled tissue was stained for EdU with a kit (C10337, Invitrogen). When done in combination with antibody staining, EdU tracing followed the secondary antibody incubation step (Hoechst was excluded from secondary antibody solution). Samples were rinsed in PBS and incubated in EdU reaction cocktail for 60 minutes at RT. Then, samples were washed in PBS, counterstained with Hoechst (1:2000 in PBS) for 20 minutes, washed in PBS, and stored in PBS before dissecting and mounting. When not in combination with antibody staining, EdU-labeled samples were rehydrated and permeabilized as above, and blocked in 3% BSA/PBS 30 minutes RT. The EdU staining was done as above, followed by washing in 3% BSA/PBS 15 minutes. The staining continued then as above.

For histology, eye samples were fixed overnight at 4°C in 4% PFA. Then, samples were rinsed in PBS, mounted in histogel, dehydrated in increasing ethanol series, and embedded in paraffin. 5-μm sections were prepared with microtome, dried overnight at 37°C and attached onto slides by brief baking at 65—70°C. The sections were stored at 4°C until use. Stainings on sections began with deparaffinization in Xylene and rehydration in decreasing ethanol series. Slides were incubated in hematoxylin for 2 minutes, rinsed in tap water, incubated in eosin for 45—60 seconds, dehydrated in increasing ethanol series and xylene, and embedded in mounting medium.

### Image acquisition

Whole mount WT, and Zebrabow samples were imaged with the Leica SP8 inverted confocal microscope, sequential scanning, with HC PL APO 10x/0.40 CS2, HC PL APO 20x/0.75 CS2, or HC PL APO 63x/1.20 CS2 objective. For Pax6 or EdU, and Hoechst staining, laser lines and emission detection ranges were 405/430—480 (Hoechst) and 488/500—550 (Pax6/EdU). For Zebrabow, laser lines and emission detection ranges were: 405/420—450 (Hoechst), 458/465— 505 (CFP), 514/521—554 (YFP), 561/585—685 (RFP). Imaris (Bitplane) was used to create optical sections and adjust signal intensity for optimal visualization of the clonal pattern of the epithelium.

Images in figures 1C and 9B were taken with Zeiss Axio Imager.M2 upright microscope, with EC Plan-Neofluoar objectives (5X/0.16, 10X/0.30). Hoechst signal was detected with excitation at 335—383 nm, and emission at 420—470 nm.

### Edu signal quantification

For EdU quantification, tile scan images with 20X objective were acquired and merged in LAS software (Leica). The amount of Edu-positive cells were determined with Imaris 9.3 (Bitplane). EdU+ cells were selected with ‘dots’ function and exported to Excel.

### Scanning electron microscopy

Fish were euthanized at selected time points by anesthesia in 0.02% Tricaine solution (Sigma) followed by decapitation. Whole heads were fixed overnight at +4°C in 2.5% glutaraldehyde (Grade 1, Sigma) in 0.1M sodium-phosphate buffer pH 7.4. On the following day, samples were rinsed in 0.1M sodium-phosphate buffer pH 7.4 three times. Eyes were dissected and stored in 0.1M sodium-phosphate buffer pH 7.4 at +4°C until further processing at the Electron Microscopy unit (University of Helsinki). Samples were treated with osmium tetroxide (2%), dehydrated in increasing ethanol series and by critical point drying. Finally, the samples were coated with platinum. Images were taken with FEI Quanta Field Emission Gun scanning electron microscope.

### Cell shape descriptor and microridge measurements

Scanning electron microscopy images were used for quantifications. Central cornea and 3—4 areas from peripheral cornea were imaged with 2000X magnification. Cell roundness and apical area were measured using ImageJ. A grid of crosses was superimposed onto the image, and cells marked by crosses, and showing clear borders, were selected with the polygon selection tool. Shape descriptors and area were measured for each cell. 3 eyes per condition were used for measurements. For microridge total and average length per cell, a subset of cells selected for shape descriptor analysis was analysed as follows. Cell was selected with the polygon selection tool, and the background was cleared. The image was then smoothened 1—3 times, brightness and contrast were adjusted automatically, and the image was convolved and converted into binary image. Finally, the pattern was skeletonized, and the ‘analyze skeleton’ function used for measuring microridge values.

### RNA-sequencing and computational analysis

For RNA-sequencing, same-age fish were grouped one day prior to the experiment into four tanks (14 fish per 3 litres). Aged matched (145, 148, 180 or 183 day-old) control and wound specimens were collected from the same tank. In respect to wounding, abrasion in both corneas of animals was performed as described above. Control animals were anesthetized and placed onto sponge for an equal duration as for wounded animals. After 1.5 hours’ recovery, fish were anesthetized again in 0.02% Tricaine and decapitated. Eyes were enucleated and cornea dissected in PBS. For one sample, 6 corneas from 3 individuals were pooled in 100 μl of Tri reagent (T9424, Sigma). RNA was isolated from TRI reagent (T9424, Sigma) using Precellys beads and standard protocols (P000912-LYSK0-A.0, Bertin Instruments). RNA was further purified using the Qiagen RNeasy MiniElute Cleanup kit (Qiagen, #74204). Polyadenylated mRNA was purified from 50 ng of Total RNA using NEBNext Poly(A) mRNA Magnetic Isolation Module (NEB, #E7490). Library preparation was completed on purified mRNA using the NEBNext Ultra Directional RNA Library prep kit (NEB, #E7420L) using 14 cycles of PCR amplification and indexed using single (i7) indexing. Indexed library preps from each sample were then pooled and sequenced at a pool concentration of 1.3 pM on the NextSeq 500 using a NextSeq High Output 75 cycle flow cell (Illumina) with 75SE reads. Basecalling and demultiplexing was performed using Illumina bcl2fastq (v2.20.0.422) software. Reads were mapped to to zebrafish GRCz11 genome using STAR aligner (2.6.0c). Gene counts were calculated using featureCounts tool from Subread package (v1.22.2) using Ensembl release 97 zebrafish gtf-files. Differential expression analysis was performed using R package DESeq2 (v1.22.2). One sample was excluded based on visual PCA inspection (Fig S2A). For downstream analyses, only genes with an entrezgene id were consider, therefore excluding predicted genes (Table S1). Gene Ontology (GO) term analysis was performed using g:Profiler (version e98_eg45_p14_ce5b097, Raudvere, et al., 2019) online tool with default parameters. Volcano plots and heatmaps were generated using GraphPad Prism 8 (version 8.3.0).

### Statistical analysis

Data were analyzed in Graphpad Prism 8 software. Data was tested for normality, and due to at least one non-normally distributed group in all data sets, non-parametric tests were used. For experiments with small n-number, non-parametric test was chosen by default. Differences between two groups were tested with Man-Whitney two-tailed test. Differences between multiple groups were tested with Kruskal-Wallis test, followed by Dunn’s multiple comparisons test. Correlations were studied with Spearman’s correlation.

## Author contributions

Conceptualization: K.I., V.S., F.M.; Methodology: K.I., V.S., F.M.; Validation: K.I., V.S., F.M.; Formal analysis: K.I., V.S., F.M.; Data analysis: K.I., V.S., F.M.; Writing: K.I., V.S., F.M.; Supervision: F.M.; Project administration: F.M.; Funding acquisition: F.M.

## Acknowledgments

The authors thank Sini Raatikainen for her technical assistance, and xxx for the critical reading of the manuscript. We also thank Pertti Panula for the access to the Zebrafish unit, and Henri Koivula for the guidance and help with the zebrafish experiments. This research was supported by the Academy of Finland, the Jane and Aatos Erkko Foundation (Jane ja Aatos Erkon Sa◻a◻tio◻).

Imaging was performed at the Electron Microscopy unit and the Light Microscopy Unit, Institute of Biotechnology, supported by HiLIFE and Biocenter Finland.

---

**Supplementary Figure 1. Clonal pattern of corneal epithelium before wounding, 3 hours post-wound, and 24 hours post-wound. A.** A maximum intensity projection of the cornea. **B, C**. Optical section (2.5 um thick) of the basal epithelial cell layer (**B**) or superficial epithelial cell layer (**C**) of central cornea pre-wound) or the wound site (3HPW, 24HPW). D. Maximum intensity projection of the nuclei of central cornea (pre-wound) or the wound site (3HPW, 24HPW). Scale bars: 200 um in A, 33 um in B,C,D.

**Supplementary Figure 2. A**. Principal component analysis of showing all samples initially collected, and the sample (wound #1) excluded from the final gene expression analysis. **B.** Gene ontology analysis (biological process) for genes with the log2 fold-change ≥1.5 or ≤ −1.5 (p-adjusted ≤0.05).

## Notes

### Competing Interest Statement

The authors have declared no competing interest.

