## Supplementary figures and images for "Unilateral corneal insult in Zebrafish results in a bilateral cell shape and identity modification, supporting wound closure"

### Figure S1

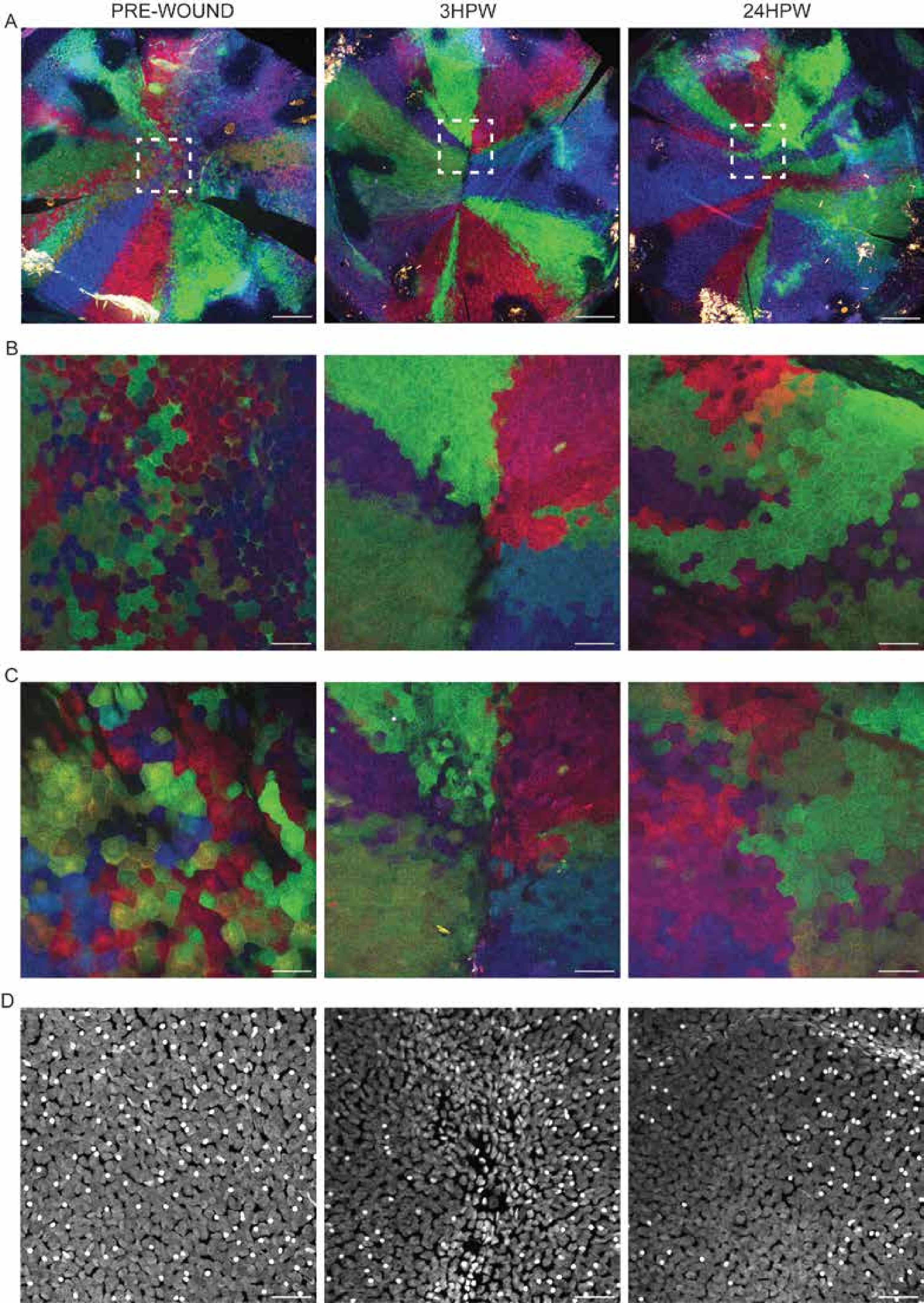

### Figure S2

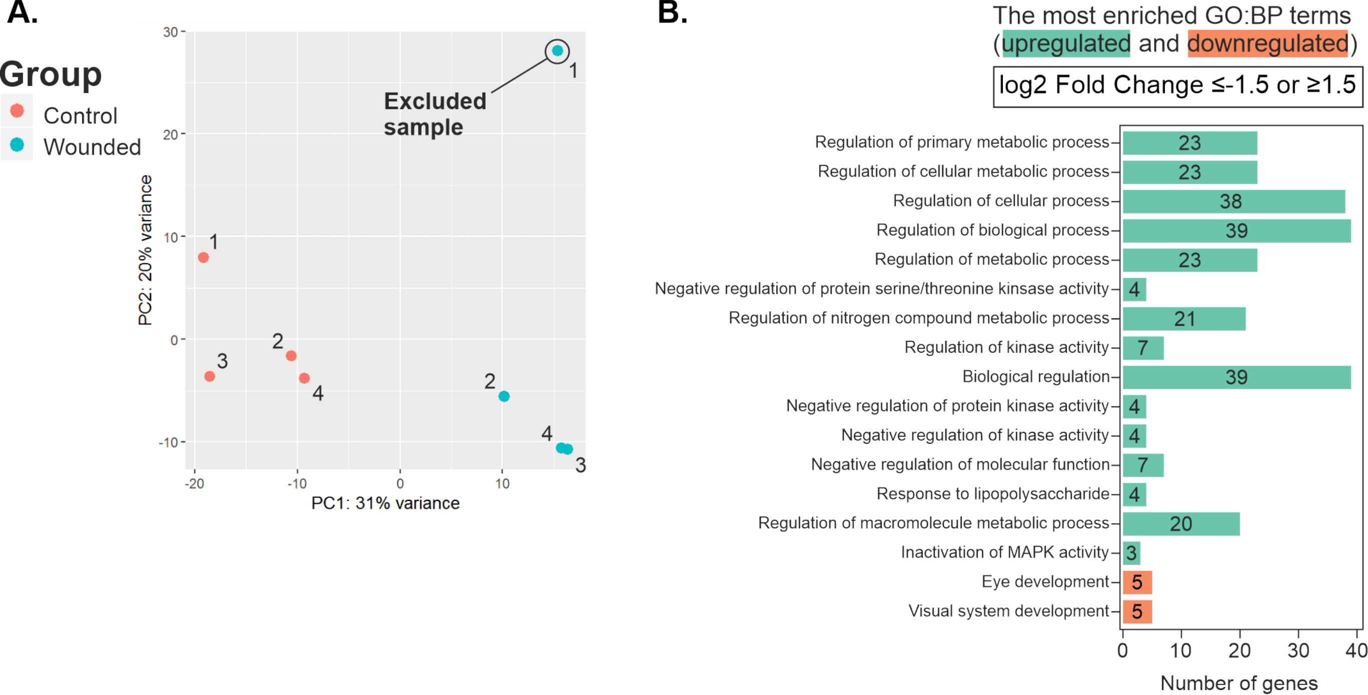
